# From partial to whole genome imputation of SARS-CoV-2 for epidemiological surveillance

**DOI:** 10.1101/2021.04.13.439668

**Authors:** Francisco M Ortuño, Carlos Loucera, Carlos S. Casimiro-Soriguer, Jose A. Lepe, Pedro Camacho Martinez, Laura Merino Diaz, Adolfo de Salazar, Natalia Chueca, Federico García, Javier Perez-Florido, Joaquin Dopazo

## Abstract

**Background:** the current SARS-CoV-2 pandemic has emphasized the utility of viral whole genome sequencing in the surveillance and control of the pathogen. An unprecedented ongoing global initiative is increasingly producing hundreds of thousands of sequences worldwide. However, the complex circumstances in which viruses are sequenced, along with the demand of urgent results, causes a high rate of incomplete and therefore useless, sequences. However, viral sequences evolve in the context of a complex phylogeny and therefore different positions along the genome are in linkage disequilibrium. Therefore, an imputation method would be able to predict missing positions from the available sequencing data.

**Results:** We developed impuSARS, an application that includes Minimac, the most widely used strategy for genomic data imputation and, taking advantage of the enormous amount of SARS-CoV-2 whole genome sequences available, a reference panel containing 239,301 sequences was built. The impuSARS application was tested in a wide range of conditions (continuous fragments, amplicons or sparse individual positions missing) showing great fidelity when reconstructing the original sequences. The impuSARS application is also able to impute whole genomes from commercial kits covering less than 20% of the genome or only from the *Spike* protein with a precision of 0.96. It also recovers the lineage with a 100% precision for almost all the lineages, even in very poorly covered genomes (< 20%)

**Conclusions:** imputation can improve the pace of SARS-CoV-2 sequencing production by recovering many incomplete or low-quality sequences that would be otherwise discarded. impuSARS can be incorporated in any primary data processing pipeline for SARS-CoV-2 whole genome sequencing.

## Background

SARS-CoV-2 is a 30 kb single stranded RNA non-fragmented virus. It is classified, together with HCoV-OC43, HCoV-HKU1, SARS-CoV-1, MERS-CoV, into the β coronaviridae. SARS-CoV-2 was first described in Wuhan, China, in December 2019, and is responsible for COVID-19, which was declared by WHO as a pandemic in March 2020 [1]. Whole genome sequencing (WGS) has been successfully used for classification [2], studying transmission dynamics [3], and evaluating global and regional patterns of pandemic spread [4]. WGS has also the potential to study reinfections, which have been described in a number of patients [5], and has very recently gained protagonism to characterize viral variants that may escape the neutralizing activity of the antibodies produced by vaccines [6]. Unfortunately, WGS results, especially in complex scenarios like this pandemic, are often imperfect, rendering incomplete viral sequences, with significant regions of the genome poorly covered [7]. Actually, current systems for viral lineage identification, a highly relevant step for the control of potentially harmful strains, fail to provide a lineage assignment if a percentage (typically > 50%) of the viral sequences is missing [8]. Given the short response times required in clinics, resequencing low-quality results is frequently not an option. Therefore, alternatives to improve sequencing results, used in other fields, such as genotype imputation, would be extremely useful in this scenario as well. Genotype imputation has traditionally been a crucial component of genome-wide association studies, by increasing the power of the findings, helping in their interpretation and facilitating further meta-analysis [9]. Genotype imputation relies on the existing correlation between genetic variants at sites across the genome of an organism [10]. Using this correlation, imputation methods accurately assign genotypes at untyped markers, improving genome coverage [10–14]. The accuracy of this imputation process improves as the number of haplotypes in the reference panel of sequenced genomes increases [15, 16], especially for variants present at low frequencies (minor allele frequency < 0.5%). The accuracy can also be increased with large reference panels. In the case of human genomes, the Haplotype Reference Consortium14, composed of about 32,000 individuals, is considered a large panel, able to reach an accurate imputation for variants with frequencies of 0.1–0.5% or less [14]. In the case of SARS-CoV-2, the outstanding international effort of sequencing has generated in a short time span a genomic database ten times larger. In spite of the interest in whole genome sequencing (WGS) viral studies and the fact that typically the sequences are imperfect, with positions and regions missing, the imputation, with a few exceptions [17], has scarcely been used in the viral realm. In addition, since WGS may not be routinely available for clinical laboratories, protocols for partial sequencing of SARS-CoV-2 genome, or even partial sequencing of the spike, where most of the determinants for variant characterization are located, are becoming available [18]. Given the importance of sequencing viral whole genomes for epidemiologic surveillance purposes, as stressed by the World Health Organization [19] and the European Parliament [20], a tool for genotype imputation in SARS-CoV-2 would increase the sequencing throughput by recovering many sequences discarded by low quality, that still contain valid information for lineage or clade assignation. Similarly, sequencing kits that only cover some key stretches already miss (or will miss future) relevant mutations. Imputation may predict the existence of these variants of interest (VOI) or variants of concern (VOC) because of their linkage disequilibrium (LD) with resolved parts of the viral genome. Here we present a fully tested, highly accurate reference panel and tool for the imputation of SARS-CoV-2 whole genome sequences from incomplete or partial sequences.

## Implementation

### SARS-CoV-2 Imputation

SARS-CoV-2 sequences’ imputation (impuSARS) was performed by using the Minimac software [14]. Although Minimac was originally designed for human samples with diploid genotypes, the tool allows imputing haploid genomes as SARS-COV-2 since it supports imputation for non-PAR regions at human males’ chromosome X. The reference panel was built with Minimac3 whereas Minimac4 was used for imputation. Minimac4 provides comparable imputation qualities as Minimac3, but it reduces memory usage and computational costs. The impuSARS tool accepts both FASTA sequence or variation (VCF) inputs. Note that FASTA sequence can include missing regions (usually tagged as N), which will be then imputed. FASTA input is aligned to reference with Muscle [21] to retrieve mutation positions. Also, VCF input should include both variant and reference genotypes when available.

The provided reference panel was created with the available SARS-CoV-2 sequences from GISAID [22, 23] (downloaded on January 7, 2021). Only sequences including >29kb and <1% missing bases were kept (“complete” and “high coverage” tags in GISAID, respectively). Also, sequences were converted to a multi-sample VCF format to only compute variant positions. As defined by GISAID, the hCoV-19/Wuhan/WIV04/2019 sequence (accession number *EPI_ISL_402124*) was considered the official reference sequence. From this multi-sample VCF, unique variants, that is, private variants for each sequence, were discarded. Therefore, the final reference panel contained 239,301 sequences. The parameter estimation for the reference panel was already precomputed with Minimac (version 3) to speed up the imputation process (reference panel provided in M3VCF format). This reference panel will be periodically updated to allow the collection of novel variants, especially VOIs and VOCs.

Once the variant imputation is performed using the previous reference panel, impuSARS will retrieve the imputed consensus sequence provided by *bcftools consensus* v1.11 [24]. Also, the associated lineage for each imputed consensus sequence will be obtained with *PANGOLIN* v1.10.2 [8]. *PANGOLIN* assigns a detailed lineage identifier to each sequence based on a multinomial logistic regression model [25]. *PANGOLIN* classifies sequences along a hierarchical tree reflecting evolutionary events. Each level of the hierarchical tree gathers a group of sequences with a common evidence associated with an epidemiological event (usually related with new variations), which could produce an emerging edge of the pandemic [25]. Lineages becoming important in the lowest levels of the phylogeny are retagged with aliases to avoid infinite spread across the hierarchical tree, thus keeping it compacted in four levels at most.

### Code availability

The imputation tool impuSARS has been encapsulated in a Docker container for interoperability and easy distribution purposes [26] and it is freely available at https://github.com/babelomics/impuSARS.

### Validation procedure

SARS-CoV-2 imputation was evaluated by using a 10-fold cross-validation process. The dataset was randomly partitioned in 10 test subsets. For each test subset, the imputation panel was computed for the remaining 9 datasets (training subsets). Initially, the loss of genomic regions was simulated by progressively increasing the percentage of the missing genome by 10% intervals. Three different strategies were used to select these missing regions: (i) random selection of only one missing region (continuous block); (ii) random selection of variant positions (missing sites) and (iii) random selection of amplicon regions that are usually independently amplified in SARS-CoV-2 sequencing (missing discontinuous blocks). Amplicon regions were defined by the hCoV-2019/nCoV-2019 v3 Amplicon Set [27] recommended by the ARTIC network [28]. Missing regions for amplicons were simulated as percentages of amplicons completely uncovered. The whole learning-testing procedure was repeated three times to reduce bias produced by the random selection. Additionally, imputation was also validated by iteratively removing a sliding window of 3kb (~10% of the entire genome) by 1,5kb steps. This process will allow determining those hotspot regions in the SARS-CoV-2 genome which are harder to impute if missed.

After validating imputation with several random selections, two more real scenarios were considered: i) imputation from regions covered by the genotyping assay kit *DeepChek*®-*8-plex CoV-2* [29]; and ii) imputation only from variants belonging to the Spike protein (S) region. As above, a 10-fold cross-validation process was implemented in both cases. The genotyping assay covers several selected regions which represent around 20% of the entire SARS-COV-2 genome, hence imputation can provide a more comprehensive, improved result. Alternatively, S protein is one of the most commonly sequenced regions for SARS-CoV-2 given its crucial role in the docking receptor recognition and cell membrane fusion [30, 31]. Moreover, mutations in spike have been related to transmissibility or the ability to evade the host immune response [32]. Therefore, studying the ability of imputing the entire SAR-CoV-2 genome from the spike region can benefit subsequent lineage classification, thus being crucial for epidemiological surveillance.

In order to facilitate the interpretation of the results we have computed the precision, recall and F1 scores. Since this is a heavily unbalanced problem (much lower number of variants against reference positions), we also provide the Matthews correlation coefficient (MCC) and Balanced accuracy (BACC) scores which are better suited for handling such scenarios [33–35]. Recall determines the true-positive rate whereas precision represents the positive predictive value. The F1-score represents the harmonic mean of the previous two metrics. The MCC measures the correlation and agreement between the truth and the predicted labels and varies between −1 and 1, where −1 refers to complete disagreement between the predicted and truth labels, 0 an average random prediction and 1 a perfect prediction. Finally, the balanced accuracy is the arithmetic mean of sensitivity and specificity.

### Lineage classification

Imputations from simulated genotyping assay and spike region test subsets were also evaluated in terms of the lineage assigned to the imputed sequences. A standard accuracy metric was calculated to evaluate assigned lineages from imputed sequences against real lineages from original GISAID sequences. Additionally, two baseline models were implemented to evaluate the influence of known variants against missing ones over the assignment of lineages. The first baseline model simply filled missing regions with the SARS-CoV-2 reference sequence. The second model randomly generated the genotype to the missing variant positions of the entire test subset weighting probabilities by the original genotype frequency in the training datasets. For comparison purposes, lineages were also obtained for the resulting sequences using these two baseline models.

### Imputation test with independent datasets

After the entire validation process, the final reference panel including the 239,301 GISAID sequences was built. Several independent datasets were considered for this test phase using the definitive reference panel: i) new GISAID sequences not included in the reference panel belonging to lineages of interest; ii) eight samples sequenced at the Hospital San Cecilio (Granada, Spain) by using both the *DeepChek*®-*8Plex-CoV2* genotyping array [29] and WGS as described below, and iii) one sample, assigned to the B.1.351 (South African lineage) [36] by an experimental RT-PCR kit, subjected to WGS that resulted in an incomplete whole-genome sequence, at Hospital Virgen del Rocio (Seville, Spain).

In the first test, new GISAID sequences from highly relevant lineages like B.1.1.7 (British lineage) [37] and B.1.351 (South African lineage) [36] were selected: 64,398 and 970 sequences, respectively (sequences downloaded by February 23rd, 2021). As in the previous validation phase, these sequences were also tested by iteratively removing a 3kb window sliding by 1,5kb steps in the entire genome. In this way the importance of specific regions to impute relevant lineages could be evaluated. In the second test the variations obtained by the genotyping array were used to impute the entire genome and the assigned lineages are compared against wholegenome results. Finally, the imputation tool was used in a third test to solve a real case in which an experimental RUO test warned of a potential VOC but the confirmatory WGS was of poor quality in a scenario where a quick informed decision was required. Then, the poor-quality sequence was used to impute the whole-genome sequence and lineage. The resolution of this case proves the level of resolution and accuracy of the imputation procedure presented here.

### RT-PCR detection of variants SARS-CoV-2 B.1.1.7, B.1.351 and B.1.1.28.1

An alternative experimental detection of variants SARS-CoV-2 B.1.1.7, B.1.351 and B.1.1.28.1, was performed by RT-PCR using a RUO kit (SARS-CoV-2 variants RT-PCR, Vitro SA) to detect the presence and/or absence of specific targets in ORF1ab gen (deletion SGF 3675-3677) and Spike gen (deletion HV 69-70).

### Genotyping array and whole genome sequencing of viral samples

Eight SARS-CoV-2 naso-pharingeal samples were sequenced following the manufacturer DeepChek®-8Plex-CoV2 genotyping array protocol [29]. WGS of the same samples was carried out following the ARTIC protocol [28] with the hCoV-2019/nCoV-2019 v3 Amplicon Set [27]. Whole-genome samples were sequenced in a NextSeq 500 sequencer by Illumina with 150bp paired-end reads and a total coverage of about 500k reads per sample.

### Sequence data preprocessing

Sequencing data (150bpx2) were analyzed using in-house scripts and the *nf-core/viralrecon* pipeline software [38]. Briefly, after read quality filtering, sequences for each sample were aligned to the SARS-CoV-2 isolate Wuhan-Hu-1 reference genome (MN908947.3) using bowtie 2 algorithm [39], followed by primer sequence removal and duplicate read marking using *iVar* [40] and *Picard* [41] tools respectively. Genomic variants are identified through *iVar* software, using a minimum allele frequency threshold of 0.25 for calling variants and a filtering step to keep variants with a minimum allele frequency threshold of 0.75. Using the set of high confidence variants and the MN908947.3 genome, a consensus genome per sample is finally built using *iVar*.

## Results

### Imputation of randomly simulated missing regions

Each of the 10 test subsets in the 10-fold cross-validation was reduced by randomly simulating missing regions in increasing percentages (10%-90%). This process was repeated 3 times for each missing percentage. Classification metrics (MCC, BACC and F1-score) were obtained for each reduced test dataset as shown in Figure 1A for one random region (missing continuous blocks), Figure 1B for randomly selected variants (missing sites) and Figure 1C for randomly selected amplicons (missing discontinuous blocks). In all cases, imputation performance metrics averaged >0.65 even for the worst scenario (imputing only with 10% of known genome). Imputation progressively improves when known sequence percentages are increasing, reaching average values >0.95 for those tests with 90% known genomes. Interestingly, the performance metrics presented a higher dispersion (including some lower outliers) when imputing only 10% of the genome in one continuous block (Figure 1A) whereas this dispersion is more marked at the opposite side of the range of values, for 90% missing regions for missing variants and discontinuous blocks (Figure 1B and C). This behavior might be related to the fact that leaving only one small random block to impute can involve regions where mutations are rare and harder to impute, even with the remaining 90% known ones. The imputation by missing sliding windows proposed below (see next Section) will help to confirm that hypothesis. Finally, even for extremely high missing percentages like the genotyping assays (~80%) or only spike regions used below, the obtained metrics suggest a reasonably accurate imputation.

**Figure 1.**
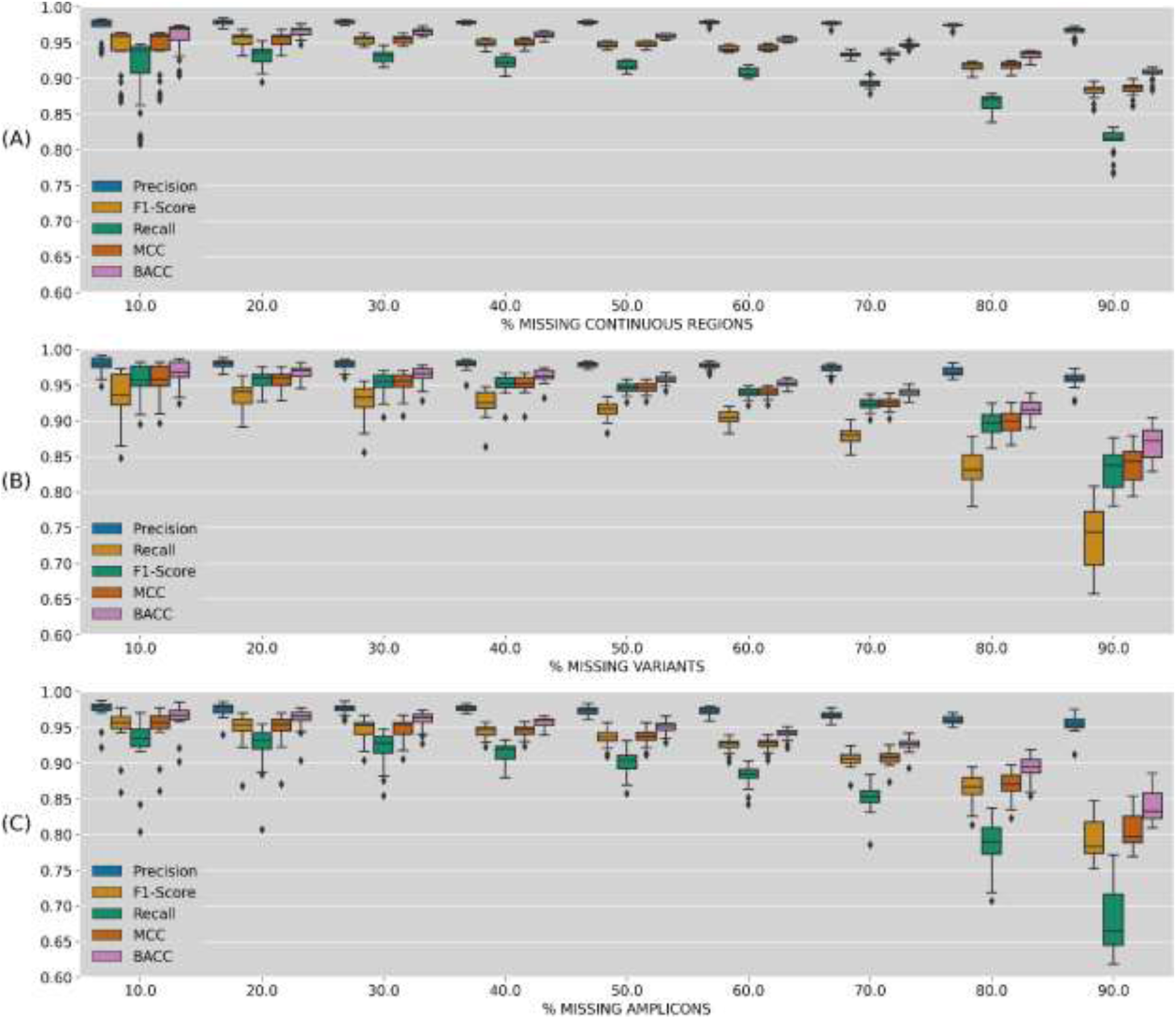
Imputation performance metrics (precision, recall, F1-score, MCC and BACC) depending on missing genome percentage. (A) One random continuous block of the genome; (B) Random selection of missing variants; (C) Random selection of missing amplicons

### Effects of missing specific locations

As previously noted, imputation performance is strongly associated with the region missing coverage in the SARS-CoV-2 genome. Therefore, the importance of selecting adequate regions when sequencing SARS-CoV-2 samples and its influence in a subsequent imputation of the remaining regions is analyzed here. For this purpose, a 3kb window was iteratively removed and imputed from the entire genome, repeating the process by 1.5kb steps. For the sake of clarity, only key metrics such as precision, recall and MCC of each imputed window along the entire genome are shown in Figure 2. Additional metrics BACC and F1-Score are available at Additional File 1: Fig.S1. Several hotspots (4 regions) have been identified as critical positions where variants are harder to impute when the block around is missing. More specifically, uncovered regions in positions around 3k, 12k, 16.5k (*orf1ab* protein, replicase polyprotein 1ab) and 24k (*S* protein, spike glycoprotein) would slightly reduce imputation ability. As previously suggested, note that those identified hotspots are strongly associated with regions where variants are less frequent in the reference panel (“*dashed green*” line).

**Figure 2.**
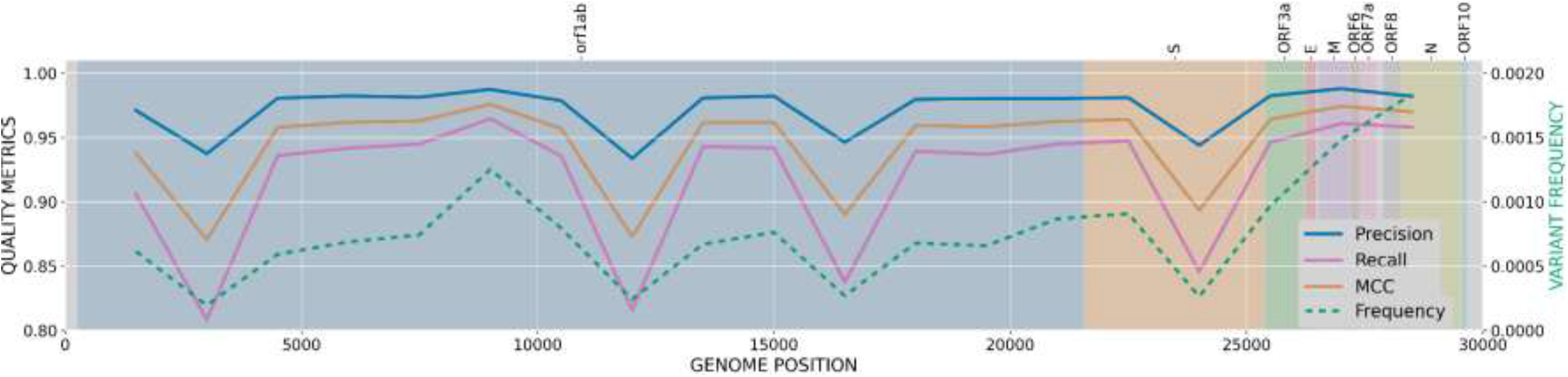
Imputation performance metrics (precision, recall and MCC) based on the position of a missing 3kb window along the SARS-CoV-2 genome. Left y-axis values represent variant frequencies (dashed green line). SARS-CoV-2 protein regions are represented by colored background and names specified at the top.

### Imputation from genotyping assay and spike regions

Once we have validated the robustness of our imputation against different missing regions scenarios, the validation will focus on the imputation of variants for sequences reduced to the genotyping assay regions previously described and Spike protein regions. Table 1 shows imputation performance metrics for both cases per test subset. Also, these metrics were calculated against the frequency of imputed variants in the reference panel (Figure 3). In both cases, we kept only the representative metrics precision, recall and MCC. Detailed results for the other mentioned metrics (BACC and F1-Score) can be found in Additional File 1: Table S1 and Additional File 1: Fig S2. As shown in Table 1, the imputation performance overcomes 0.81 in the three averaged metrics, being precision the highest with >0.96 for both regions while recall remains at 0.86 and 0.81 for genotyping assay and spike regions, respectively. Regarding Figure 3, variant imputation quickly raises to >0.96 in the three performance metrics (recall, precision and MCC) for variants with frequencies >0.01 and >0.03 for the genotyping array and spike region imputations, respectively. The imputation from genotyping array sequences reaches its maximum values (>0.996) from frequencies over 0.33 for precision and recall metrics, whereas MCC slightly drops to 0.895 after the same frequency threshold. For imputation from the spike region, an improvement is also observed from variant frequencies >0.33 reaching performance values of 0.998 and 0.969 for recall and precision, respectively but a more drastic fall is observed in MCC. This MCC decrease is correlated in both cases with the drop in the number of variants (“green” line). When variant frequency increases, a smaller number of variants are found but datasets are inversely unbalanced (more variant than reference positions) which metric-wise is better captured by the MCC. Nevertheless, imputing positive cases (variants) in those situations are more relevant, so results in recall and precision metrics are more informative.

**Table 1.** Performance metrics (Recall, Precision and MCC)

| Subset | Imputation from genotyping assay kit |  |  | Imputation from Spike region |  |  |
| --- | --- | --- | --- | --- | --- | --- |
|  | Recall | Precision | MCC | Recall | Precision | MCC |
| 1 | 0.8595 | 0.9612 | 0.9088 | 0.8129 | 0.9618 | 0.8841 |
| 2 | 0.8578 | 0.9597 | 0.9072 | 0.8121 | 0.9620 | 0.8838 |
| 3 | 0.8562 | 0.9614 | 0.9072 | 0.8100 | 0.9625 | 0.8829 |
| 4 | 0.8609 | 0.9622 | 0.9101 | 0.8106 | 0.9616 | 0.8828 |
| 5 | 0.8589 | 0.9603 | 0.9081 | 0.8109 | 0.9619 | 0.8831 |
| 6 | 0.8593 | 0.9602 | 0.9083 | 0.8106 | 0.9608 | 0.8824 |
| 7 | 0.8586 | 0.9600 | 0.9078 | 0.8126 | 0.9613 | 0.8837 |
| 8 | 0.8597 | 0.9614 | 0.9091 | 0.8106 | 0.9624 | 0.8831 |
| 9 | 0.8579 | 0.9605 | 0.9077 | 0.8115 | 0.9622 | 0.8835 |
| 10 | 0.8574 | 0.9609 | 0.9076 | 0.8121 | 0.9629 | 0.8842 |
| Avg±Std | 0.8586<br>±0.0013 | 0.9608<br>±0.0008 | 0.9082<br>±0.0009 | 0.8114<br>±0.0010 | 0.9619<br>±0.0006 | 0.8834<br>±0.0006 |

Metrics obtained for 10-fold cross-validation subsets imputing from the genotyping assay and Spike protein regions. Values are calculated for the entire test subset imputation.

**Figure 3.**
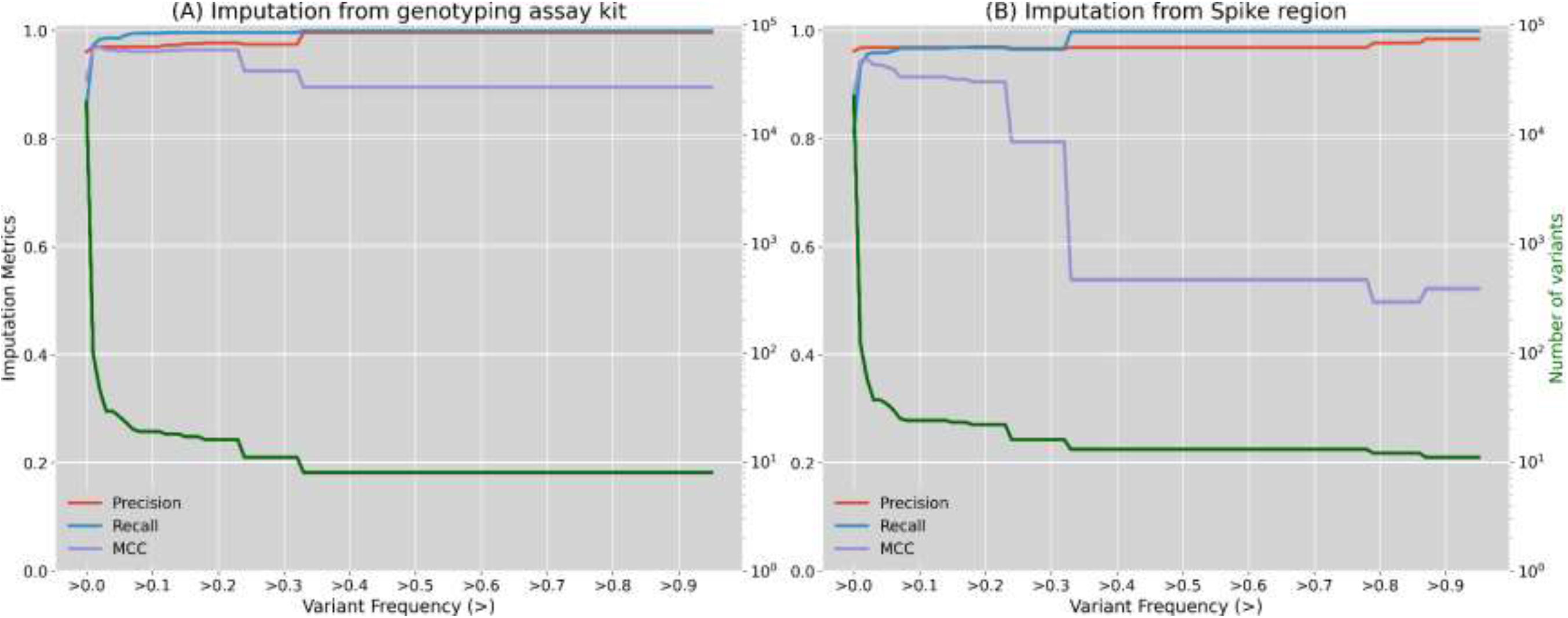
Principal imputation performance metrics (precision, recall and MCC) calculated depending on imputed variant frequencies. (A) Imputation quality when imputing from the genotyping array positions; (B) Imputation quality when imputing from spike protein positions. Left y-axis (green) represents the number of variants for those frequency threshold (log scale)

### Lineage classification

The previously imputed variants for the simulated genotyping arrays and spike region subsets are used to rebuild the consensus whole genome sequences and assign their corresponding lineages with PANGOLIN. The quality of the imputed lineage has been measured by the accuracy metric against real lineages and compared to two baseline models (Figure 4). Briefly, these two models respectively filled missing regions with random variants assigned by frequency (“Random fill”) or with nucleotides from the reference sequence (“Reference fill”) (see Implementation section for details). Also, accuracy was calculated for the different levels of the hierarchical tree in PANGOLIN lineages. As shown, the first level in the hierarchical classification of lineage was almost always correctly determined (>98%), even for the two baseline models. That is, the information provided by the already known regions (genotyping array and spike protein) was enough to classify this first level. However, the imputed solution becomes more relevant as a lower level has to be determined. Hence, imputation clearly outperformed both baseline methods when lineages were assigned at 3rd and 4th level, achieving 77% and 68% accuracy for genotyping array and spike regions, respectively. As expected, imputation from the genotyping array positions comes up with higher lineage accuracies than the solution with spike, since this kit was specifically designed to capture relevant regions in the SARS-CoV-2 genome. Even so, imputation still produces strong benefits in the lineage assignment for the genotyping array regions, clearly improving lineage assignments with simple baseline models.

**Figure 4.**
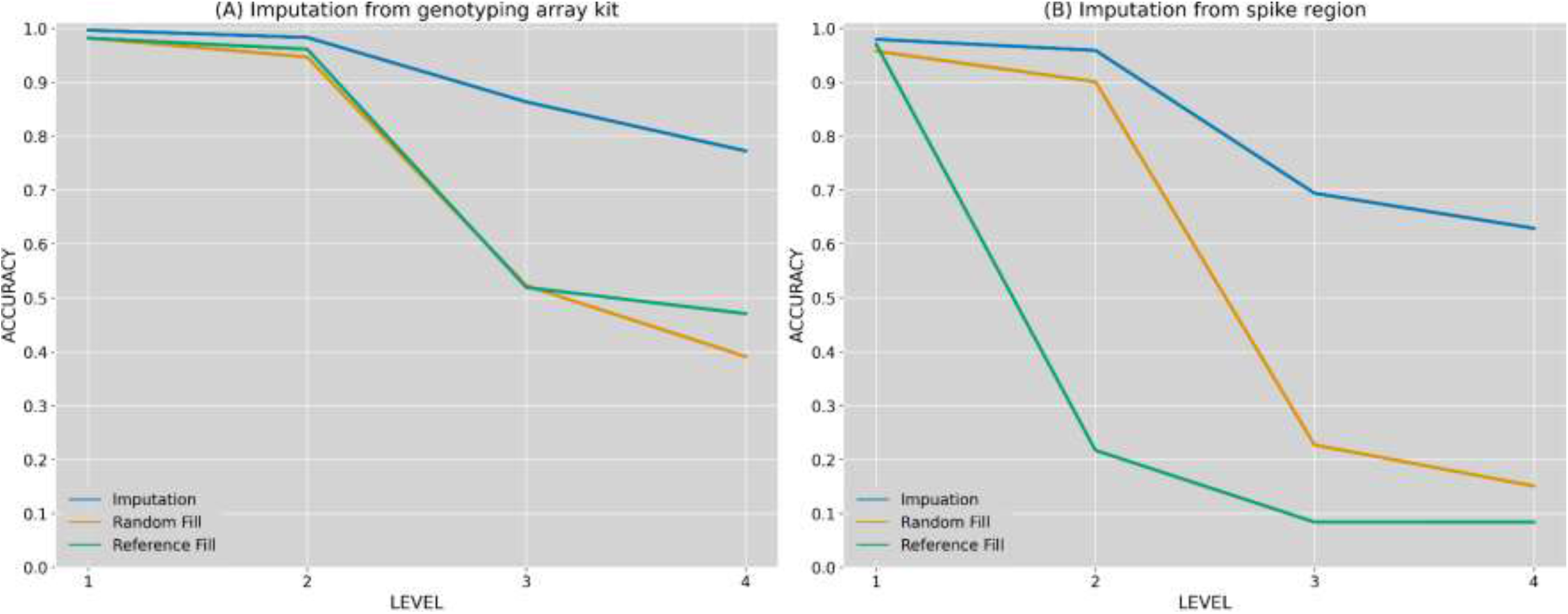
Lineage classification accuracy compared against two baseline models. (A) Lineage accuracy when imputing from the genotyping array positions; (B) Lineage accuracy when imputing from spike protein region. Levels represent lineage specification.

Additionally, a detailed view about lineage classification for the top frequent lineages (>500 sequences) is shown in Figure 5. As noted, there are lineages that are more commonly misclassified. For instance, several sequences are wrongly classified as B.1.1.119 when imputing from the genotyping array regions. Similarly, lineage B.1 is frequently assigned when sequences truly belong to a more specific lineage (lower level in the hierarchical tree) in the imputation from spike. In the first case, this misclassification is produced by the fact that lineage B.1.1.119 is partially constituted by three variants in positions 28881-28883, which are not captured by the used genotyping array. This situation makes sequences from other close lineages like B.1, B.1.1.214 or B.1.1.282 identical to B.1.1.119, from the genotyping array perspective. Consequently, these close lineages are frequently imputed as B.1.1.119 (30%, 88% and 73%, respectively). Likewise, given the lack of certain regions when imputing from spike region, several sub-branches like B.1.1.119, B.1.1.214, B.1.1.282, or B.1.1.284 are wrongly classified as the parent node B.1 (80%, 75%, 87% and 57%, respectively).

**Figure 5.**
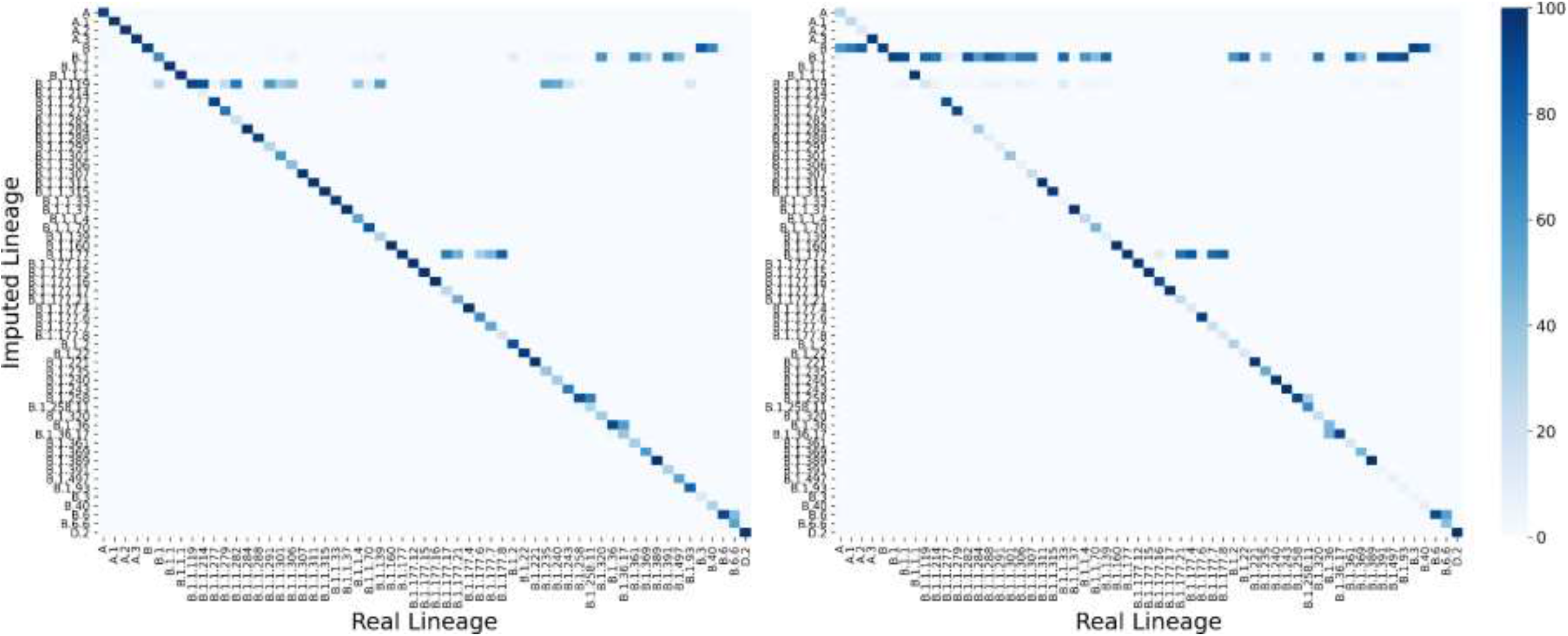
Accuracy obtained for each pair of lineages (real vs imputed) for the top frequent lineages (>500 sequences). Left heatmap represents the obtained values for genotyping array imputation whereas right heatmap represents accuracies for imputation from spike protein region. Color represents the percentage of sequences in each real lineage classified by each imputed lineage (the darker, the higher).

### Imputation of new independent datasets

Previous sections have extensively validated the proposed imputation system under several configurations and strategies. This section will show several use cases and test results produced by independent datasets over the final imputation reference panel (239,301 sequences).

Firstly, two recently emerging lineages, B.1.1.7 (British lineage) and B.1.351 (South African lineage), have also been studied in this final testing phase to evaluate the performance of the imputation in new lineages. Sequences recently added to GISAID (not included in our current reference panel) under these lineages were selected: 64,398 and 970 sequences, respectively. Their percentage of correctly classified lineages after imputation when missing a 3kb window (10%) along the entire genome are then calculated (Figure 6).

**Figure 6.**
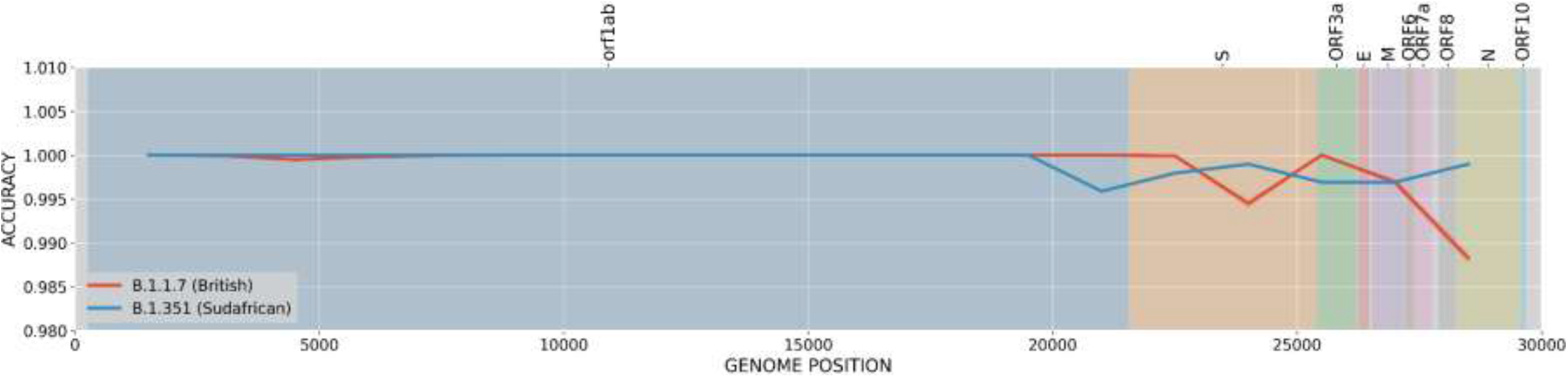
Lineage classification accuracy. Accuracy is estimated for a missed region in sliding windows of 3kb for the recent British and South African lineages (B.1.1.7 and B.1.351, respectively)

As shown in Figure 6, even when these lineages are underrepresented in the current reference panel (23 and 105 sequences, respectively), the methodology has captured the LD structure at such precision that it can accurately impute the B.1.1.7 and B.1.351 lineages from other sequences. Specifically, both lineages obtained 100% accuracy for almost any missing 3kb region. The imputation accuracy resulted slightly reduced in the British lineage (B.1.1.7) when the missing regions are located around the center of S protein (99.5% accuracy) or at ORF8 and N proteins (99% accuracy). This behavior is clearly associated with the loss of constitutive variants for the British lineage like N501Y, A570D or P681H, among others [42]. In the case of the South African lineage (B.1.351), performance vaguely dropped at the beginning of protein *S* (99.5%) as well as around *E* and *M* proteins (99.8%). Again, these small decreases are associated with important variants associated with the lineage like Q57H or P71L [43].

### Imputation for sequencing kits and low-quality sequences

Eight SARS-CoV-2 samples were sequenced using the *DeepChek*®-*8-plex CoV-2* genotyping array (see Table 2). The partial sequences covering about 20% of the whole viral genome were used to impute the remaining non-covered 80% genome with impuSARS. Then, the same samples were subjected to WGS. The imputed whole genome sequences and lineages were subsequently compared against each other, rendering a highly reliable imputation sequence and 100% successful lineage imputation. FASTQ files as well as consensus whole genome sequences for both genotyping array and whole-genome sequencing of these 8 samples are available for download at the European Nucleotide Archive (ENA) under the accession ID PRJEB43882. Also, imputation results (both imputed consensus whole genome sequences and lineages) are provided in a Zenodo repository (https://doi.org/10.5281/zenodo.4616731). Coverage distribution from initial genotyping array results are provided in Additional File 1: Fig. S3. The three main quality metrics and imputed lineages are shown in Table 2. A more detailed table including variant counts and additional metrics is provided in Additional File 1: Table S2.

**Table 2.** Variant imputation metrics (precision, recall and MCC) and lineage classification

| Sample | Recall | Precision | MCC | Real Lineage | Imputed |
| --- | --- | --- | --- | --- | --- |
| AND00023 | 0.9000 | 1 | 0.9486 | B.1.1.7 | B.1.1.7 |
| AND00040 | 0.8571 | 1 | 0.9258 | B.1.1.7 | B.1.1.7 |
| AND00065 | 0.8636 | 1 | 0.9293 | B.1.1.7 | B.1.1.7 |
| AND00073 | 0.8571 | 1 | 0.9258 | B.1.1.7 | B.1.1.7 |
| AND00123 | 0.9231 | 1 | 0.9607 | B.1.1.7 | B.1.1.7 |
| AND00128 | 0.6000 | 1 | 0.7745 | B.1.1.7 | B.1.1.7 |
| AND00132 | 0.8696 | 1 | 0.9324 | B.1.1.7 | B.1.1.7 |
| AND00139 | 0.9091 | 1 | 0.9534 | B.1.1.7 | B.1.1.7 |
| Avg ±Std Dev | 0.8475±0.103 | 1.0000±0.00 | 0.9188±0.06 | | 100% |

Values for eight independent samples internally sequenced with both the genotyping array and whole-genome sequencing.

To further illustrate the usefulness of the imputation system in a real clinical scenario, a use case of the Hospital Virgen del Rocio is described. In a routine survey a sample was analyzed by RT-PCR using a RUO kit (see Implementation section for details), which raised a warning suggesting it may belong to the emerging South African lineage (B.1.351), a VOC. The sample was immediately submitted to confirmatory WGS, that resulted in a poor-quality sequencing, with only 28.91% of SARS-CoV-2 genome covered, having 71 amplicons completely non-covered and 3 covered at low depth (<20x). Lineage assignation with current tools like PANGOLIN is impossible in this low-quality scenario. However, there was an urgency in confirming or discarding the presence of a VOC for epidemiologic surveillance and medical decision making. Therefore, impuSARS was used on this poor-quality sequence and lineage imputation was carried out with PANGOLIN producing a B.1.1.7 lineage assignment, also a VOC, but currently more extended in Spain. Detailed analysis of the pattern of available mutations also supported this lineage assignation (See Table 3).

**Table 3.** Study of AND00344 variants

| Mutation | Found in variant | Present | Coverage | South Africa | UK | Brazil |
| --- | --- | --- | --- | --- | --- | --- |
| L18F | South Africa / Brazil | no | none | ? |  | ? |
| T20N | Brazil | no | none |  |  | ? |
| P26S | Brazil | no | none |  |  | ? |
| del_21765 | UK | no | none |  | ? |  |
| D80A | South Africa | no | none | ? |  |  |
| D138Y | Brazil | no | none |  |  | ? |
| del_21991 | UK | no | none |  | ? |  |
| R190S | Brazil | no | none |  |  | ? |
| D215G | South Africa | no | none | ? |  |  |
| del_22281 | South Africa | no | covered | no |  |  |
| R246I | South Africa | no | covered | no |  |  |
| K417N | South Africa / Brazil | no | none | ? |  | ? |
| E484K | South Africa / Brazil | no | low | no? |  | no? |
| N501Y | UK / South Africa / Brazil | yes | covered | yes? | yes? | yes? |
| A570D | UK | no | none |  | ? |  |
| D614G | UK / South Africa / Brazil | no | none | ? | ? | ? |
| H655Y | Brazil | no | covered |  |  | no |
| P681H | UK | yes | covered | no | yes | no |
| A701V | South Africa | no | covered | no |  |  |
| T716I | UK | yes | covered | no | yes | no |
| S982A | UK | no | none |  | ? |  |
| T1027I | Brazil | no | none |  |  | ? |
| D1118H | UK | no | none |  | ? |  |
| Q57H | South Africa | no | covered | no |  |  |
| P71L | South Africa | no | covered | no |  |  |
| Q27stop | UK | yes | covered | no | yes | no |
| T205I | South Africa | no | low | no? |  |  |
|  |  |  |  | NO | YES (Most likely) | NO (most likely) |

Comparison of the available variation in the low coverage sequence of vial sample AND00344 with respect to the South Africa (B.1.351), UK (B.1.1.7) and Brazil (P.1) VOCs

## Conclusions

Whole genome sequence imputation from partial sequences from commercial kits or from low-quality WGS has demonstrated to produce highly reliable results and be an excellent tool for lineage assignment. Given the short response times required for the identification of samples for decision support or for epidemiological surveillance in a clinical context, re-sampling and/or resequencing are not realistic options. Therefore, imputation constitutes an accurate and useful tool to complement and improve SARS-CoV-2 WGS pipelines in clinics.

## Availability and requirements

**Project name:** impuSARS (SARS-CoV-2 imputation)

**Project home page:** https://github.com/babelomics/impuSARS

**Operating system(s):** Platform independent (Docker container)

**Programming language:** Python, Bash

**Other requirements:** Docker

**License:** MIT License.

**Any restrictions to use by non-academics:** none

## List of abbreviations

BACC: Balanced accuracy
LD: linkage disequilibrium
MCC: Matthews correlation coefficient
RT-PCR: Real Time Polymerase Chain Reaction
RUO: Research use only
VCF: Variant Calling Format
VOC: variants of concern
VOI: variants of interest
WGS: Whole Genome Sequencing

## Declarations

### Ethics approval and consent to participate

Not applicable

### Consent for publication

Not applicable

### Availability of data and materials

- The SARS-CoV-2 sequences used to train the impuSARS tool were taken from GISAID: https://www.gisaid.org/epiflu-applications/hcov-19-reference-sequence/
- The hCoV-19/Wuhan/WIV04/2019 sequence (EPI_ISL_402124) was taken from GISAID: https://platform.gisaid.org/epi3/start/CoV2020
- The 8 SARS-CoV-2 whole genome sequences generated in this study are available at the European Nucleotide Archive: https://www.ebi.ac.uk/ena/browser/view/PRJEB43882.
- The Imputation results (both imputed whole genome sequences and lineages) are provided in the Zenodo repository: https://doi.org/10.5281/zenodo.4616731

### Authors’ contributions

FO performed most of the analysis and wrote the draft of the manuscript, CL carried out the statistic part of the work, CCS and JPF contributed to the analysis of the samples, JAL, PCM, LMD carried out the use case of the RUO kit, AS, NC and FG carried out the commercial kit use case, and JD conceived the work and wrote the manuscript

## Additional Files

**Additional file 1**

PDF format .PDF

Supplementary Tables and figures:

**Table S1. Supplementary imputation performance metrics (BACC and F1)**

**Table S2. Variant counts and additional metrics**

**Fig. S1. More imputation performance metrics (F1 and BACC) based on the position of a missing 3kb window along the SARS-CoV-2 genome**.

Left y-axis values represent variant frequencies (dashed green line). SARS-CoV-2 protein regions are represented by colored background and names specified at the top.

**Fig. S2. Supplementary imputation performance metrics (BACC and F1) calculated depending on imputed variant frequencies.** (A) Imputation quality when imputing from the genotyping array positions; (B) Imputation quality when imputing from spike protein positions. Left y-axis (green) represents the number of variants for those frequency threshold (log scale)

**Fig S3. Coverage distribution from genotyping array in the eight samples studied.**

**Fig. S1. More imputation performance metrics (F1 and BACC) based on the position of a missing 3kb window along the SARS-CoV-2 genome**.

Left y-axis values represent variant frequencies (dashed green line). SARS-CoV-2 protein regions are represented by colored background and names specified at the top.

