## Additional file 1 for "From partial to whole genome imputation of SARS-CoV-2 for epidemiological surveillance"

### Imputation of SARS-CoV-2 whole genome sequences from incomplete or partial sequences

#### Supplementary Tables:

**Table S1. Supplementary imputation performance metrics (BACC and F1)**

**Table S2. Variant counts and additional metrics**

#### Supplementary Figures:

**Fig. S1. More imputation performance metrics (F1 and BACC) based on the position of a missing 3kb window along the SARS-CoV-2 genome**.
Left y-axis values represent variant frequencies (dashed green line). SARS-CoV-2 protein regions are represented by colored background and names specified at the top.

**Fig. S2. Supplementary imputation performance metrics (BACC and F1) calculated depending on imputed variant frequencies.** (A) Imputation quality when imputing from the genotyping array positions; (B) Imputation quality when imputing from spike protein positions. Left y-axis (green) represents the number of variants for those frequency threshold (log scale)

**Fig S3. Coverage distribution from genotyping array in the eight samples studied.**

**Table S1. Supplementary imputation performance metrics (BACC and F1)**

| Subset | Imputation from genotyping assay kit | | Imputation from Spike region | |
|---|---|---|---|---|
| | BACC | F1-Score | BACC | F1-Score |
| 1 | 0,9297 | 0,9075 | 0,9064 | 0,8811 |
| 2 | 0,9289 | 0,9059 | 0,9060 | 0,8807 |
| 3 | 0,9281 | 0,9058 | 0,9050 | 0,8797 |
| 4 | 0,9305 | 0,9088 | 0,9053 | 0,8797 |
| 5 | 0,9294 | 0,9068 | 0,9054 | 0,8800 |
| 6 | 0,9296 | 0,9069 | 0,9053 | 0,8793 |
| 7 | 0,9293 | 0,9065 | 0,9063 | 0,8807 |
| 8 | 0,9298 | 0,9077 | 0,9053 | 0,8800 |
| 9 | 0,9289 | 0,9063 | 0,9057 | 0,8804 |
| 10 | 0,9287 | 0,9062 | 0,9060 | 0,8811 |
| **Average** | 0,9293 | 0,9068 | 0,9293 | 0,9068 |
| **Std Dev** | 0,0007 | 0,0009 | 0,0007 | 0,0009 |

**Table S2. Variant counts and additional metrics**

| Sample | Number of Ref Positions | | Number of Variants | | | | Recall | Precision | MCC | BACC | F1-Score |
|---|---|---|---|---|---|---|---|---|---|---|---|
| | In kit | From WGS not in kit (N) | In kit | From WGS not in kit (P) | Imputed (TP + FP) | Correctly Imputed (TP) | | | | | |
| AND00023 | 7005 | 22762 | 11 | 20 | 18 | 18 | 0,9000 | 1 | 0,9486 | 0,9500 | 0,9474 |
| AND00040 | 7313 | 22455 | 20 | 14 | 12 | 12 | 0,8571 | 1 | 0,9258 | 0,9286 | 0,9231 |
| AND00065 | 6509 | 23257 | 12 | 22 | 19 | 19 | 0,8636 | 1 | 0,9293 | 0,9318 | 0,9268 |
| AND00073 | 7297 | 22466 | 23 | 14 | 12 | 12 | 0,8571 | 1 | 0,9258 | 0,9286 | 0,9231 |
| AND00123 | 7379 | 22388 | 20 | 13 | 12 | 12 | 0,9231 | 1 | 0,9607 | 0,9615 | 0,9600 |
| AND00128 | 7448 | 22319 | 11 | 20 | 12 | 12 | 0,6000 | 1 | 0,7745 | 0,8000 | 0,7500 |
| AND00132 | 6159 | 23601 | 19 | 23 | 20 | 20 | 0,8696 | 1 | 0,9324 | 0,9348 | 0,9302 |
| AND00139 | 6425 | 23338 | 12 | 22 | 20 | 20 | 0,9091 | 1 | 0,9534 | 0,9545 | 0,9524 |
| | | | | | | **Avg** | 0,8475 | 1,0000 | 0,9188 | 0,9237 | 0,9141 |
| | | | | | | **Std Dev** | 0,1031 | 0,0000 | 0,0599 | 0,0516 | 0,0678 |

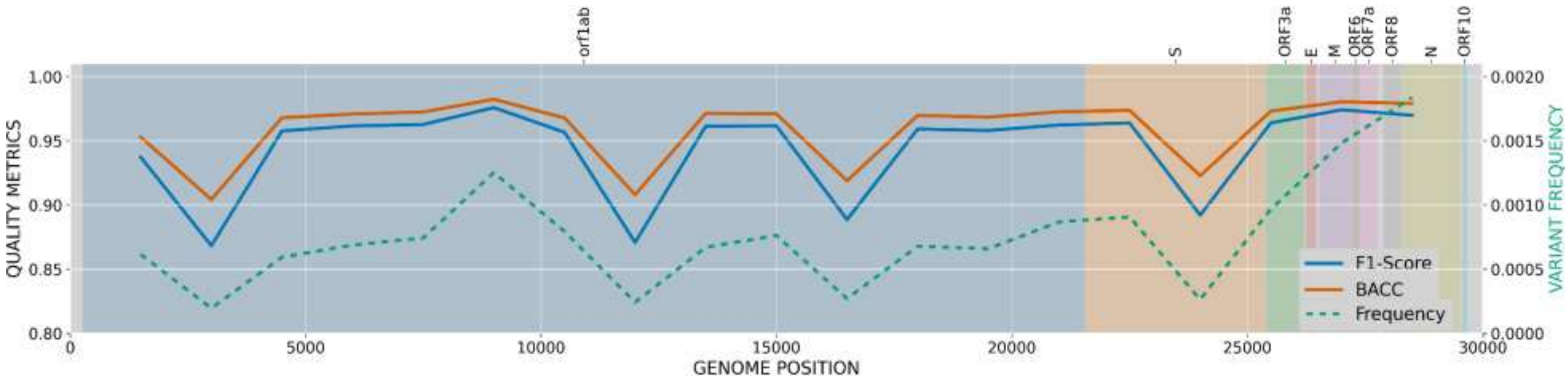


**Fig. S1. More imputation performance metrics (F1 and BACC) based on the position of a missing 3kb window along the SARS-CoV-2 genome**.

Left y-axis values represent variant frequencies (dashed green line). SARS-CoV-2 protein regions are represented by colored background and names specified at the top.

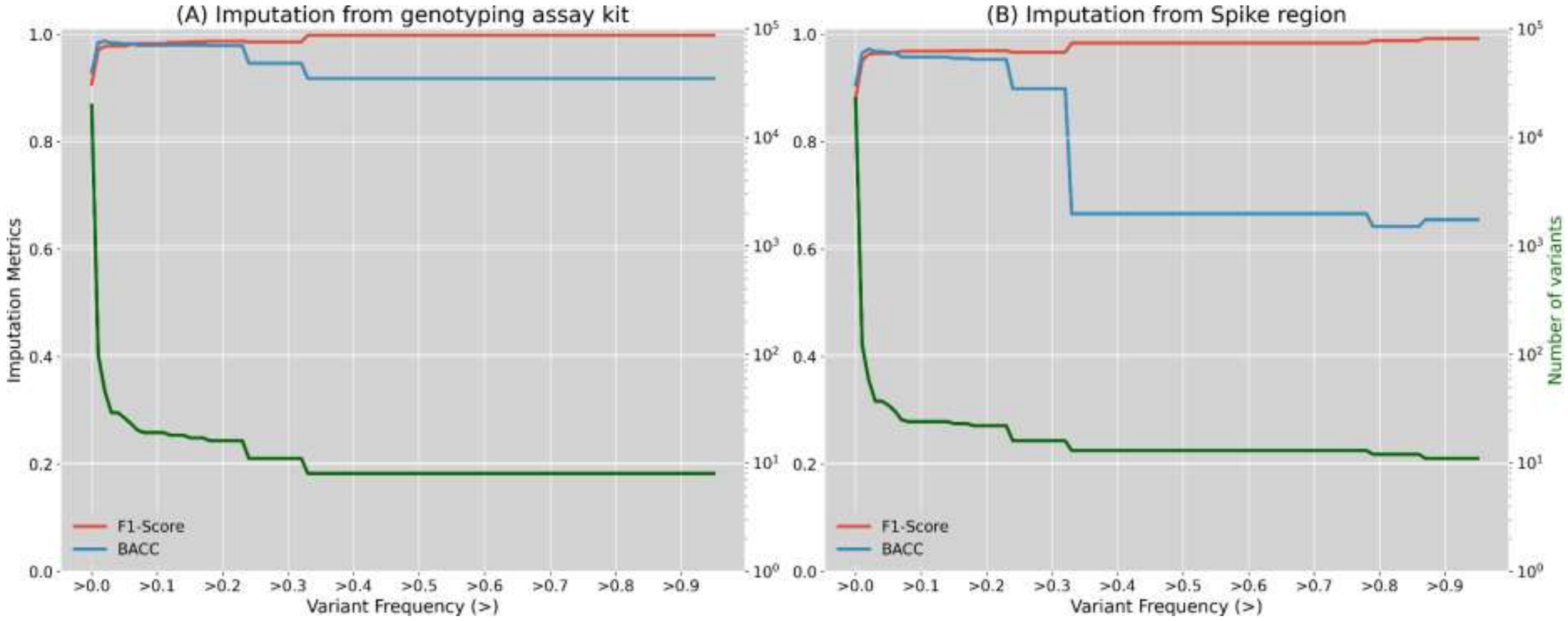


**Fig. S2. Supplementary imputation performance metrics (BACC and F1) calculated depending on imputed variant frequencies.** (A) Imputation quality when imputing from the genotyping array positions; (B) Imputation quality when imputing from spike protein positions. Left y-axis (green) represents the number of variants for those frequency threshold (log scale)

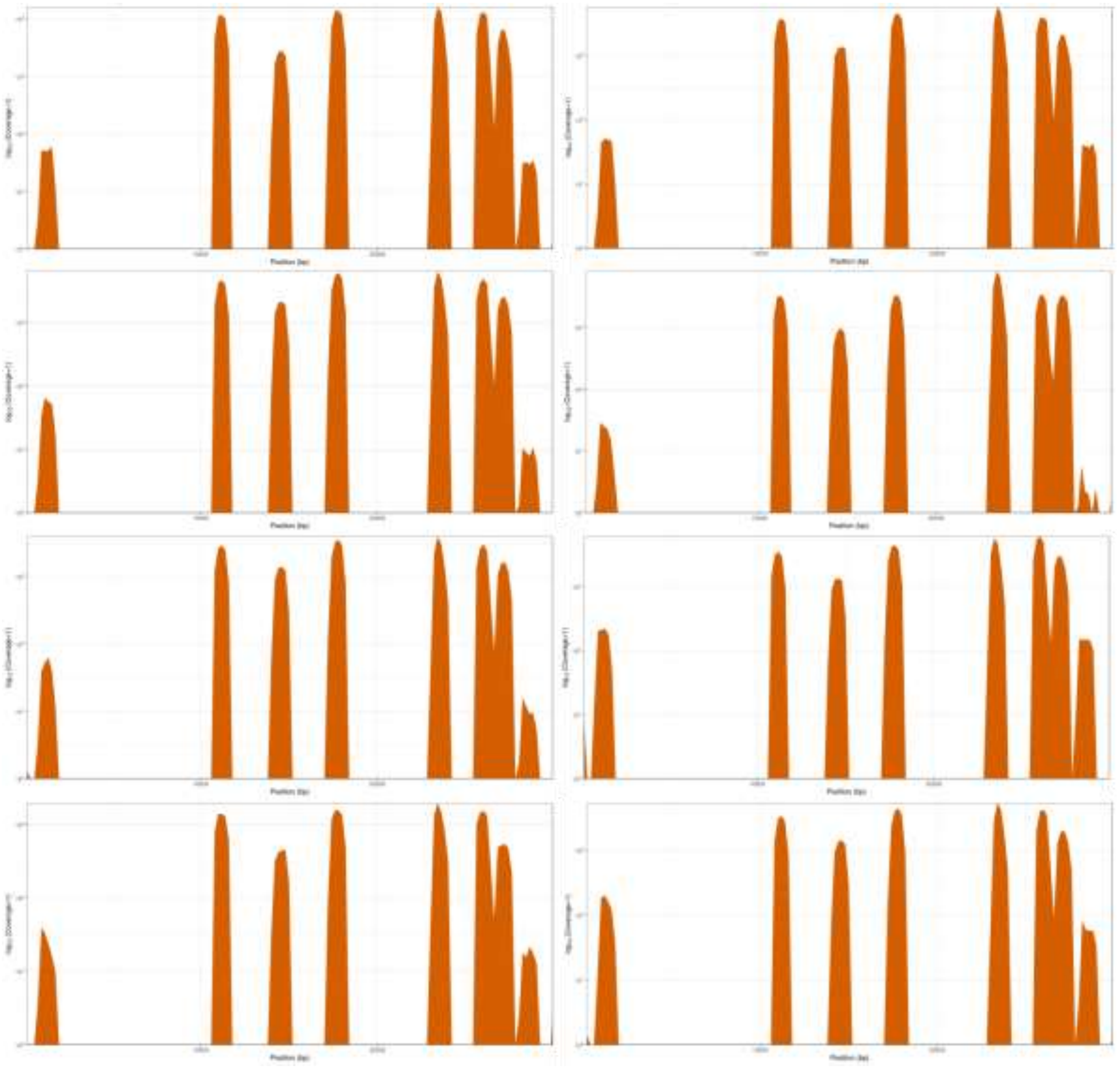

**Fig S3. Coverage distribution from genotyping array in the eight samples studied.**
